# Mitochondrial fission is required for proper nucleoid distribution within mitochondrial networks

**DOI:** 10.1101/2021.03.17.435804

**Authors:** Hema Saranya Ilamathi, Mathieu Ouellet, Rasha Sabouny, Justine Desrochers-Goyette, Matthew A. Lines, Gerald Pfeffer, Timothy E. Shutt, Marc Germain

**Affiliations:** Groupe de Recherche en Signalisation Cellulaire and Département de Biologie Médicale, Université du Québec à Trois-Rivières, Trois-Rivières, Québec, Canada; Centre d’Excellence en Recherche sur les Maladies Orphelines - Fondation Courtois, Université du Québec à Montréal, Montréal, Québec, Canada; Réseau Intersectoriel de Recherche en Santé de l’Université du Québec (RISUQ); Department of Medical Genetics, Cumming School of Medicine, University of Calgary, Calgary, Alberta, Canada; Department of Clinical Neurosciences, Cumming School of Medicine, University of Calgary; Hotchkiss Brain Institute, Department of Clinical Neurosciences, Cumming School of Medicine, University of Calgary, Calgary, Alberta; Alberta Children’s Hospital Research Institute, Department of Medical Genetics, Cumming School of Medicine, University of Calgary, Calgary, Alberta, Canada; Department of Biochemistry & Molecular Biology, Cumming School of Medicine, University of Calgary, Calgary, Alberta, Canada

**Keywords:** mitochondrial DNA, mitochondrial fission, nucleoid distribution, algorithm, DRP1, MYH14

## Abstract

Mitochondrial DNA (mtDNA) maintenance is essential to sustain a functionally healthy population of mitochondria within cells. Proper mtDNA replication and distribution within mitochondrial networks are essential to maintain mitochondrial homeostasis. However, the fundamental basis of mtDNA segregation and distribution within mitochondrial networks is still unclear. To address these questions, we developed an algorithm, Mitomate tracker to unravel the global distribution of nucleoids within mitochondria. Using this tool, we decipher the semi-regular spacing of nucleoids across mitochondrial networks. Furthermore, we show that mitochondrial fission actively regulates mtDNA distribution by controlling the distribution of nucleoids within mitochondrial networks. Specifically, we found that primary cells bearing disease-associated mutations in the fission proteins DRP1 and MYH14 show altered nucleoid distribution, and acute enrichment of enlarged nucleoids near the nucleus. Further analysis suggests that the altered nucleoid distribution observed in the fission mutants is the result of both changes in network structure and nucleoid density. Thus, our study provides novel insights into the role of mitochondria fission in nucleoid distribution and the understanding of diseases caused by fission defects.

**Significance statement:** Mutation or deletion of mitochondrial DNA (mtDNA) is associated with a large number of human diseases. However, the mechanisms controlling mtDNA replication and segregation are still poorly understood. Here, we have developed a new computational method to quantify the distribution of nucleoids (mtDNA with associated proteins) and define how nucleoid distribution is affected by changes in mitochondrial network structure. We demonstrate that mitochondrial fission is required for the proper distribution of nucleoids across mitochondrial networks, cells from patients with fission defects showing irregular nucleoid distribution and perinuclear accumulation. Nonetheless, each fission mutant behaved in a distinct manner, indicating a complex relationship between mitochondrial dynamics and nucleoid distribution.

## Introduction

Mitochondria require proteins encoded by both nuclear and mitochondrial DNA (mtDNA) to perform their key roles in cellular metabolism. Maintenance of mtDNA copy number and integrity, as well as mtDNA distribution across mitochondrial networks, is crucial for proper mitochondrial function. mtDNA is packed into nucleoprotein complexes called nucleoids. Nucleoids are dynamic structures that actively move within mitochondrial networks and interact with neighboring nucleoids (1). However, the mechanisms regulating nucleoid maintenance and distribution within mitochondrial networks are still not fully understood.

mtDNA replication is associated with mitochondrial dynamics, the process of mitochondrial fission and fusion (2). Mitochondrial fusion is regulated by the GTPases Mitofusins (MFN1 and MFN2; outer membrane) and Optic Atropy-1 (OPA1; inner membrane) and is required for maintenance of mtDNA copy number and integrity (3–5). Mitochondrial fission is mediated by dynamin related protein 1 (DRP1). DRP1-dependent fission occurs at endoplasmic reticulum (ER)-mitochondrial contact sites (ERMCS) following the initial constriction of the mitochondrial tubule by actin and myosin (6, 7). mtDNA replication occurs at ERMCS (8). Mutation of the fission proteins non-muscle myosin II (MYH14) or silencing/genetic ablation of DRP1 causes a decrease in nucleoid number, generally without a change in total mtDNA content (9–12). In the case of DRP1 deletion, these nucleoids are enlarged and confined to an abnormal modified mitochondrial structure called mito-bulbs (9, 10, 12, 13). While these results suggest that mitochondrial fission is required for nucleoid segregation, it remains unclear how fission contributes to nucleoid maintenance and the spatial distribution of nucleoids within mitochondrial networks.

A number of studies have previously measured nucleoid distribution using custom scripts (14–17). Most of these studies reported either the overall nucleoid density or the average distance between the two closest nucleoids (nearest neighbor distance; nndist), sometimes calculated without considering the positional constraints imposed by the mitochondrial network (14, 15). While these studies give an overview of the general distribution of nucleoids, other descriptors of nucleoid distribution within networks could provide a more informative description of nucleoid distribution. As such, the pair correlation function (pcf) calculates the probability of finding a nucleoid at any distance from a first one within the mitochondrial network, allowing for a finer mapping of nucleoid distribution compared with nndist (which only provides an average distance between adjacent nucleoids).

Here, we have developed Mitomate Tracker, an automated tool that evaluates the distribution of nucleoids within mitochondrial networks using both nndist and pcf. Using this tool, we demonstrate that nucleoids are distributed in a semi regular fashion within mitochondrial networks, maintaining a minimal spacing between each other. Furthermore, nucleoid density is higher in perinuclear as opposed to peripheral mitochondrial network clusters. These features were affected by mutations in MYH14 or DRP1, indicating that mitochondrial fission plays an important role in nucleoid distribution and maintenance.

## Results

### Mitomate tracker, a new tool to analyze the distribution of nucleoid across mitochondrial networks

Using the DNA binding dye picogreen, nucleoids can be visualised as punctate structures along TMRM-labeled mitochondria in live cells (Fig. 1). To study nucleoid distribution in an automated manner, we developed an algorithm (Mitomate tracker) that takes advantage of our previously-published mitochondrial network quantification tool (Momito,(18)), to which we combined two other tools: the ImageJ plugin, Trackmate which allows nucleoid identification (19), and the R package, spatstat which calculates point pattern distributions (20). This results in two distinct outputs: (1) a quantification of network and nucleoid parameters (nucleoid number and density, network length and connectivity), and (2) a point pattern distribution calculated by two distance-based metrics. The first metric, nndist, measures the average distance between a nucleoid and its closest neighbour within the mitochondrial network. The second metric, pcf, estimates the probability of finding a nucleoid at any distance from a first nucleoid within the mitochondrial network (Fig. 1).

**Figure 1:**
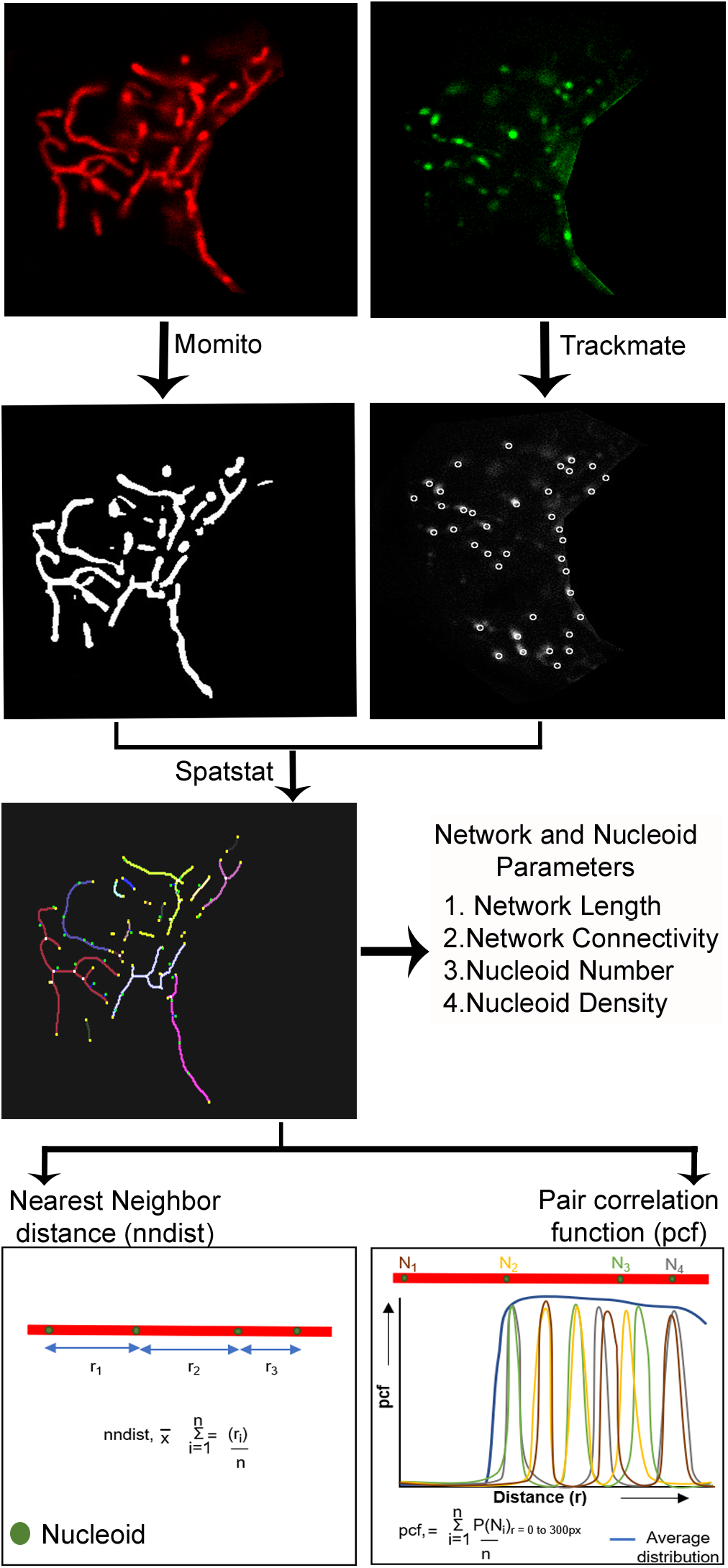
Schematic representation of nucleoid distribution analysis by Mitomate tracker. Confocal live cell Images of mitochondria (TMRM, Red) and nucleoids (Picogreen, Green – nucleus manually removed) are analyzed using Mitomate tracker. Mitochondria are segmented and their components are identified using Momito while nucleoids are identified using the Image J Plugin Trackmate. The information extracted from Momito and Trackmate is then analyzed by the R package spatstat. Mitomate Tracker provides detailed descriptors of network and nucleoid features and measures nucleoid distribution pattern by two metrics, the nearest neighbor distance (nndist) and the pair correlation function (pcf).

### Validating the robustness of nucleoid distribution metrics

As previous studies of nucleoid distribution used the average distance between adjacent nucleoids (nndist) as their primary metric (14, 16, 17), we first examined the nndist output of Mitomate tracker. Not surprisingly, the absolute distance between nucleoid was dependent on nucleoid density (Fig. 2A; r^2^=0.55, p<0.001). To take this into account, we normalised the actual nndist to an independent random process (IRP), where the same number of points are distributed within the same network independently of each other and network structure. The resulting nndist Ratio should be equal to 1 for a random distribution (actual and random values being equal). However, the normalised nndist was still somewhat dependent on nucleoid density (Fig. 2B; r^2^=0.30, p<0.001). This suggests that nndist is not a robust approach to measure nucleoid distribution owing to its persistent dependence on point density.

**Figure 2:**
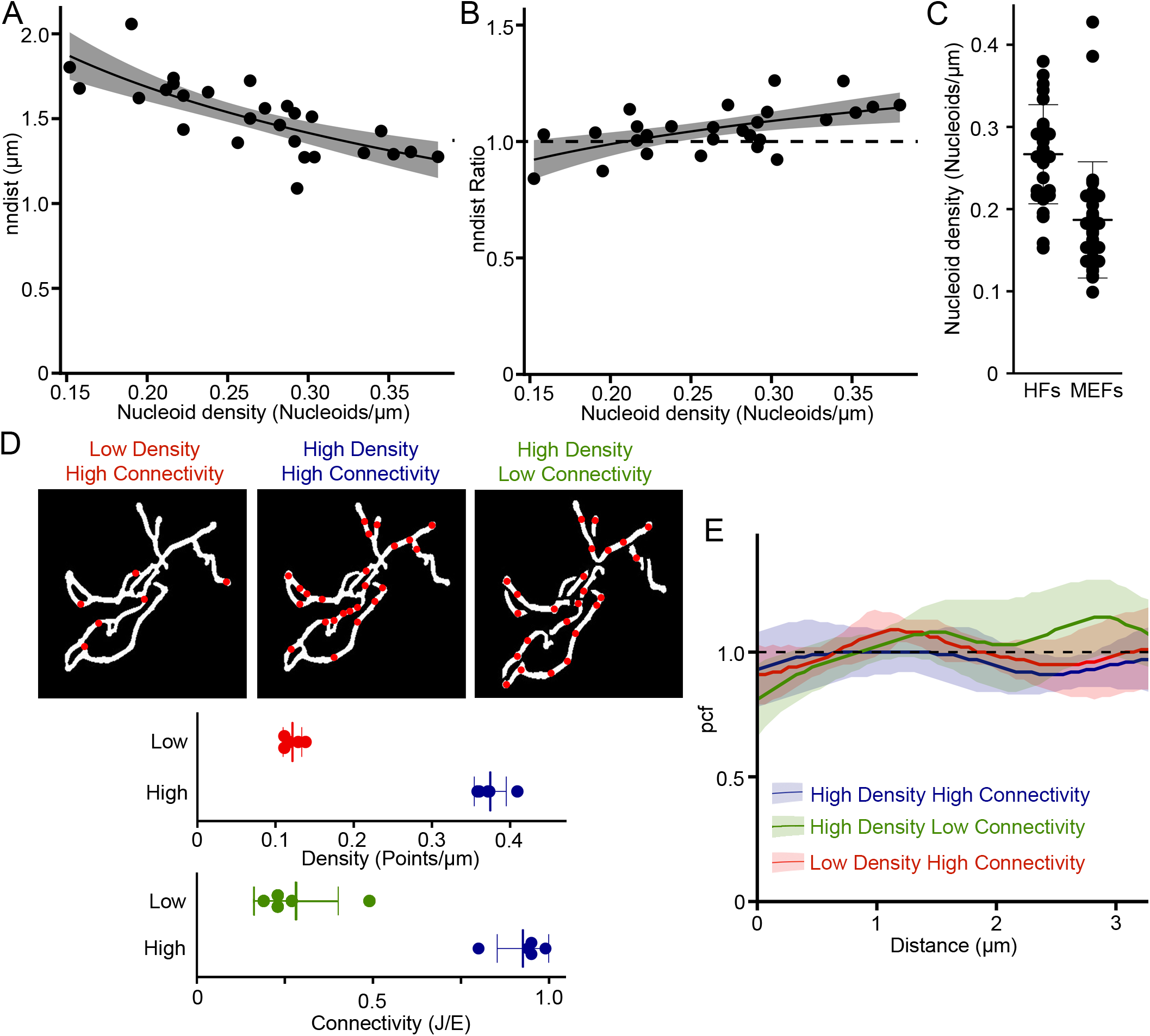
Measurement of nucleoid distribution by Mitomate tracker. (A)(B) nndist is dependent on point density even after normalization. Scatter plot of Actual (A) and normalized (relative to an IRP; B) nndist values relative to nucleoid density in primary human fibroblasts. Each data point represents an individual cell (n=27). The dashed line represents a random distribution. Linear regression formula y~log(x). (C) Nucleoid density in primary human fibroblast (HF) and mouse embryonic fibroblast (MEF) cells. Each point represents an individual cell. (D)(E) Network connectivity and point density do not affect the pcf of random point distribution. (D) The networks were either left connected or manually unbranched and overlayed with a low or a high point density. Top, Representative test network used for the analysis. Bottom, measures of point density and network connectivity for the 5 test networks. Each point represents a distinct network. (E) pcf curves. Solid lines represent the average point distribution for the indicated conditions and the shaded areas, the SD (n=5 images). The dashed line represents the expected random distribution.

We then evaluated the robustness of the pcf. Contrary to the nndist which gives a single average distance per cell, the pcf computes the probability to find a point at any distance of a first point, making it impossible to achieve a simple correlation analysis as used for nndist. We thus used a small number of highly connected mitochondrial networks to which we randomly added points representing the highest and the lowest nucleoid densities found in mouse embryonic fibroblasts (MEFs) and primary human fibroblasts (HFs) (Fig. 2C-D). To avoid measuring effects due to specific random distributions, six different distributions were averaged for each image/condition (see Methods for details on the normalisation process). Using these, we then evaluated the influence of point density on the pcf. In this analysis, a random distribution (IRP) has a pcf value of 1 at all distances from the first point (Fig 2E, black dashed line), values above 1 indicate correlation and values below 1, avoidance (Fig. S1). Consistent with this, points randomly distributed across our test mitochondrial networks resulted in pcf values close to 1 for all densities tested (Fig 2E; lines, distribution; colored area, SD). Pcf values also remained close to 1 when connectivity was reduced in our test images by manually unbranching the networks within the images using ImageJ (Fig. 2D-E), while keeping point density constant. Overall, our data indicate that the pcf provides a robust approach to study nucleoid distribution within mitochondrial networks.

### Nucleoids have a well-defined organization within mitochondrial networks

To study nucleoid distribution, MEFs and primary human fibroblasts were stained for mitochondria (TMRM) and nucleoids (picogreen) and imaged by confocal microscopy. The images were then processed and analyzed by Mitomate tracker, which identifies individual picogreen foci as a distinct nucleoid, irrespective of its mtDNA content (larger nucleoids probably containing more mtDNA copies). This allows us to analyse nucleoid distribution independently of nucleoid segregation following mtDNA replication.

The pcf showed a strong nucleoid avoidance at short distances (pcf value <1) but not at distances greater than 1 μm (pcf value >1) for both MEFs (Fig. 3A) and primary human fibroblasts (Fig. 3B). The maximal likelihood to find a neighboring nucleoid as estimated from the pcf curve peak for each cell, occurred at 1-3 μm (average 2 μm; Fig. 3C). To quantify the differences between the observed nucleoid distribution pattern and an IRP-based distribution, we then measured the entropy (Shannon entropy) of individual pcf curves. In information theory, entropy represents the amount of information present in a variable (21, 22). In the context of a pcf, this means that any horizontal line, including an IRP (pcf = 1) has an entropy of zero, and entropy increases as it deviates from this horizontal line (Fig. S1). The entropy thus provides a measure of the variability of the pcf curve. Accordingly, the entropy values of actual MEFs pcf was significantly higher than for their corresponding IRPs (Fig. 3D). Overall, our results indicate that nucleoid distribution is regulated to maintain a minimal distance between nucleoids, consistent with previous quantifications of inter-nucleoid distances using nndist in yeast cells (16, 17).

**Figure 3:**
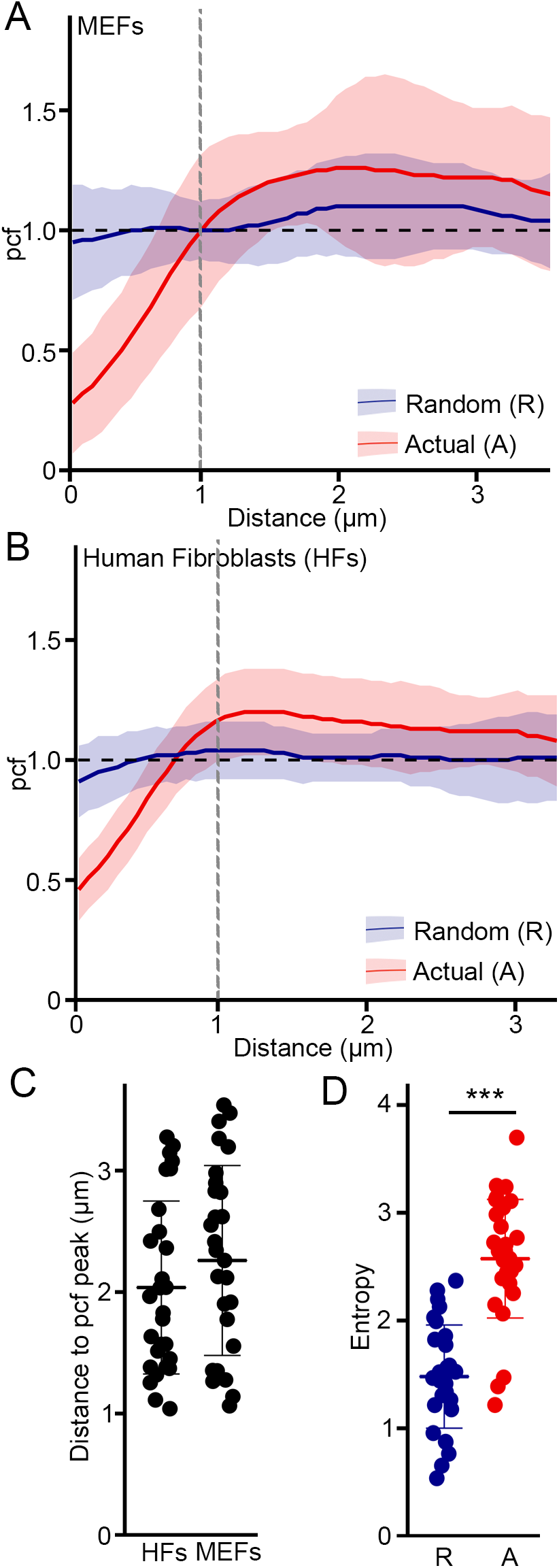
Nucleoid distribution is highly regulated. Pcf curves of nucleoid distribution in MEFs (A) and HFs(B), Solid lines represent the average of nucleoid distribution and the shaded areas the SD (n=30 cells). Actual (A, Red) and Random (R, Blue) distributions for the same point densities are shown. (C) Distance between nucleoids as determined by the distance of the maximal pcf value. (D) Entropy values calculated from MEFs pcf curves in (A). Actual nucleoid distribution (A), random distribution (R). Each data point represents one cell. Bars represent the average of 30 cells ± SD. ***p<0.001. Two-tailed t-test.

### Distribution of nucleoids within mitochondrial networks is independent of its unique spatial distribution across the cellular region

Previous studies have suggested that perinuclear mitochondria could be functionally and structurally distinct from mitochondria closer to the plasma membrane (peripheral mitochondria) (23–25). We therefore determined the spatial distribution of nucleoids relative to the nucleus in human fibroblasts. Consistent with the presence of distinct mitochondrial regions, most nucleoids were present close to the nucleus (Fig. 4A). In addition, manually separating mitochondrial clusters close to the nucleus (perinuclear (N)) from the rest of the mitochondrial network (cell periphery (P)) (Fig. 4B) revealed that nucleoid density (Number of nucleoids/ mitochondrial area) was significantly higher in the perinuclear region than in the cell periphery (Fig. 4C). Overall, both measures indicate that nucleoids accumulate close to the nucleus. We then determined whether this difference in nucleoid density affected the pcf. Perinuclear and cell periphery pcfs were almost identical to those of whole cells (Fig. 4D) resulting in similar entropy values (Fig. 4E). This indicates that these two cellular regions have a similar nucleoid distribution pattern irrespective of nucleoid density. Altogether, our data indicates that nucleoids adopt a clear spatial organization with most of them close to the nucleus, but that the nucleoid distribution pattern (low probability to find a nucleoid at short distances relative to an IRP of the same density) is independent of this organization.

**Figure 4:**
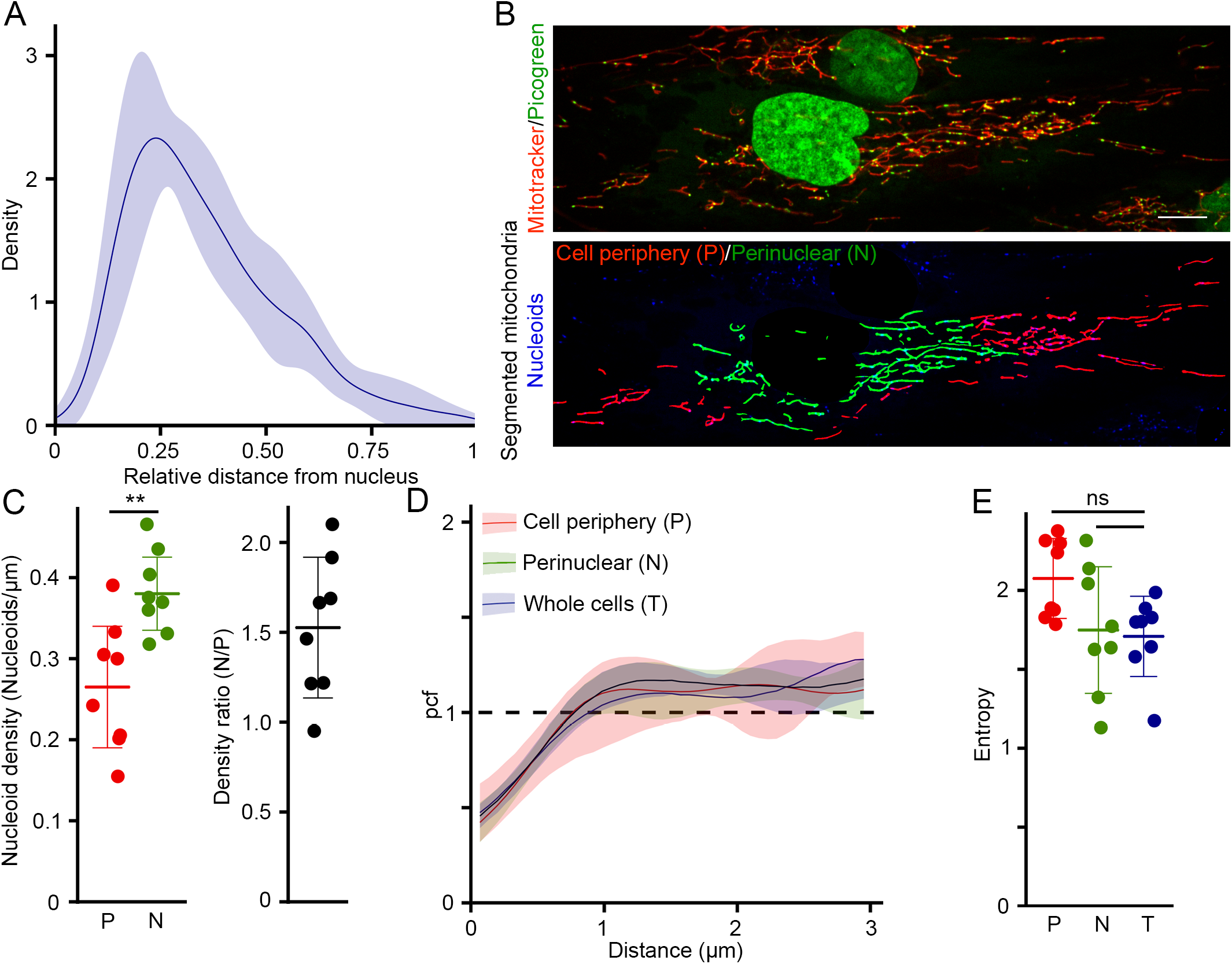
Higher nucleoid density in the perinuclear region. (A) Density plot of nucleoids from human fibroblasts relative to the nucleus, where 0 corresponds to the nucleus and 1, the cell membrane. The solid line represents the average distribution and the shaded areas the SD (n=8 cells). (B-C) Quantification of nucleoid density in the perinuclear and peripheral regions. Mitochondrial network of human fibroblasts (B) (Top; Mitochondria stained with mitotracker Red) were manually separated into peripheral (P, Red) and perinuclear (N, Green) regions (Bottom). Nucleoids were stained with picogreen (Top: Green, Bottom: Blue with nucleus removed). (C) Nucleoid density and density ratio (Perinuclear/periphery). Each data point represents one cell. Bars represent the average of 8 cells ± SD. **p<0.01. Two-tailed t-test. (D-E) Nucleoid distribution is similar irrespective of cellular localization. (D) pcf curves. Solid lines represent the average distribution for the indicated conditions and the shaded areas the SD (n=8 cells). The dashed line represents the expected random distribution. (E) Entropy calculated from the curves in (D). Each data point represents one cell. Bars represent the average of 8 cells ± SD. ns not significant. Two-tailed t-test.

### Loss of mitochondrial fission impairs the distribution of nucleoids within mitochondrial networks

Mitochondrial networks are shaped by mitochondrial dynamics, including mitochondrial fusion and fission. Importantly, sites of mitochondrial fission have been associated with sites of mtDNA replication (8). However, whether this actively contributes to nucleoid distribution within mitochondrial networks and in relation to the nucleus remains unclear. To determine the role of fission in nucleoid distribution, we investigated nucleoid content and distribution in patient fibroblasts with a dominant-negative mutation in MYH14 (R941L), a protein required for the initial constriction of mitochondrial tubules prior to DRP1-dependent scission (10). MYH14 mutation causes an increase in mitochondrial length (10) while somewhat decreasing overall mitochondrial content, but did not strongly impact mitochondrial connectivity (Fig. 5A). On the other hand, as we previously reported (10), MYH14 mutant fibroblasts had fewer nucleoids, resulting in a lower overall nucleoid density (Fig. 5A).

**Figure 5:**
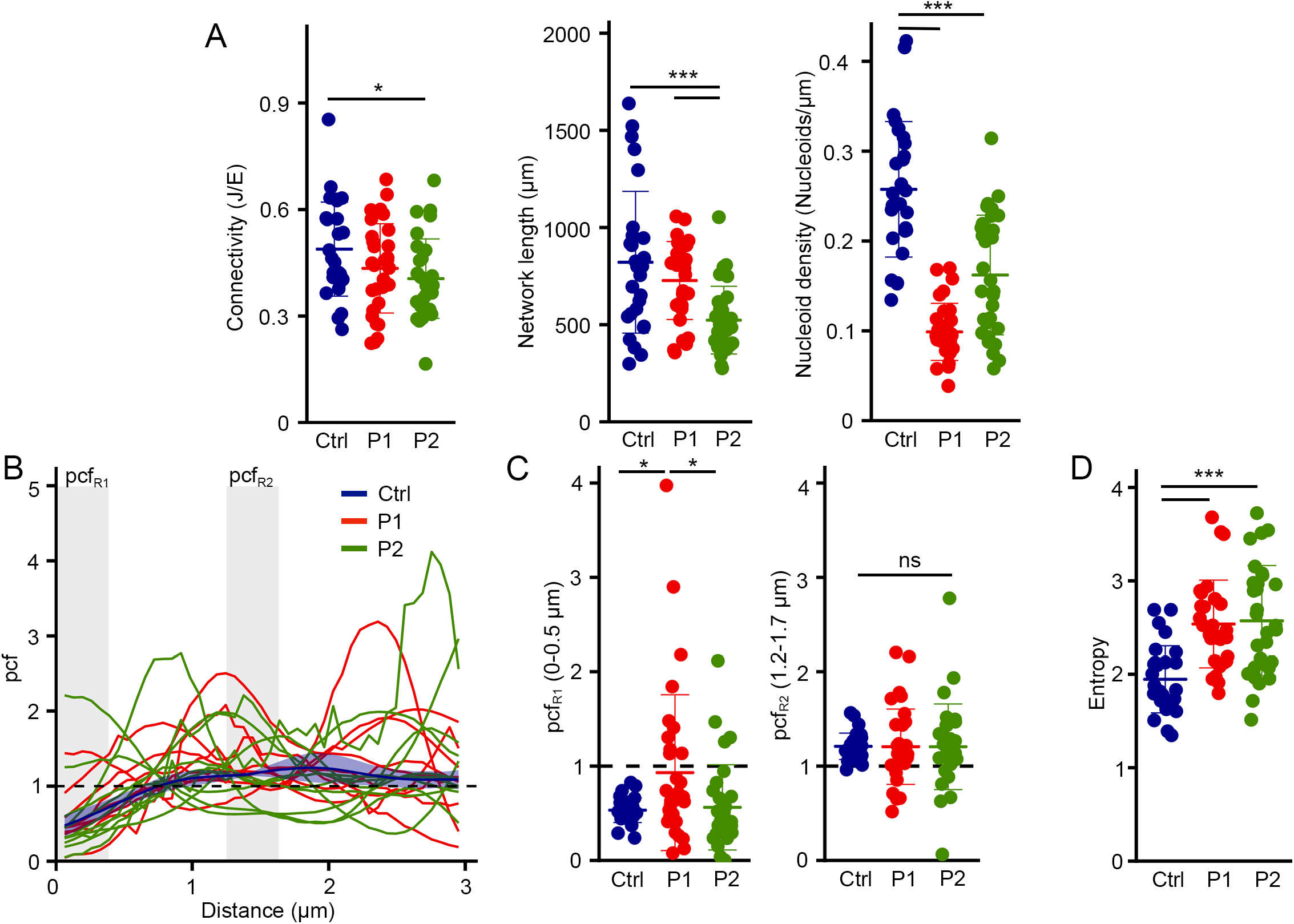
MYH14 is required for proper nucleoid distribution. (A) Mitochondrial parameters in Control and MYH14 mutant primary human fibroblasts (P1: Patient 1, P2: Patient 2). Each data point represents one cell. Bars represent the average of 30 cells ± SD. *p<0.05, ***p<0.001. One-way ANOVA. (B-D) Nucleoid distribution in control and MYH14 primary human fibroblasts. (B) pcf curves. The solid line for the control (Blue) represents the average distribution and the shaded area, the SD (n=8 cells). For the MYH14 mutants, each solid line represents one individual cell (n=9 cells per patient line), highlighting the variability of the pcf. The dashed line represents the expected random distribution and the grey areas, the distances for which the average pcf was quantified in (C; pcfR1: 0-0.5μm, pcfR2: 1.2-1.7μm). (C) Average pcf values at short and longer distances. Each data point represents one cell. Bars represent the average of 30 cells ± SD. *p<0.05, ns not significant. One-way ANOVA (D) Entropy values calculated from the pcf curves in (B). Each data point represents one cell. Bars represent the average of 30 cells ± SD. ***p<0.001. One-way ANOVA.

We then analysed nucleoid distribution in MYH14 mutants. Consistent with MYH14 mutation affecting nucleoid distribution, all mutant cells showed an altered pcf relative to control cells (Fig. 5B, each curve represents a distinct cell). However, we could not observe any conserved pattern across cells, and there were no specific changes in correlation at either short (pcfR1, 0-0.5 μm) or longer distances (pcfR2, 1.2-1.7μm) (Fig. 5B-C). In fact, while some cells showed nucleoid clustering at short distances, other cells had a strong avoidance at short distances and a distinct correlation at longer distances (Fig. 5B-C). The increased pcf variability found in MYH14 mutants was also reflected in their increased entropy (Fig.5D). Overall, the increased variance we observed in the pcf suggests that nucleoids are disorganised in MYH14 mutants in comparison to the control.

We then determined whether MYH14 mutation also alters the spatial organization of nucleoids relative to the nucleus. As with control cells, MYH14 mutants had a large fraction of their nucleoids close to the nucleus (Fig. 6A). However, the perinuclear accumulation was more pronounced in mutant cells, with nucleoids further depleted from the cell periphery (KS test, p<0.001) (Fig. 6A). The perinuclear accumulation of nucleoids in MYH14 mutant cells was further confirmed by the significant increase in density ratio (nucleoid density in the perinuclear region relative to that of the periphery) observed in mutant cells (Fig. 6B).

**Figure 6:**
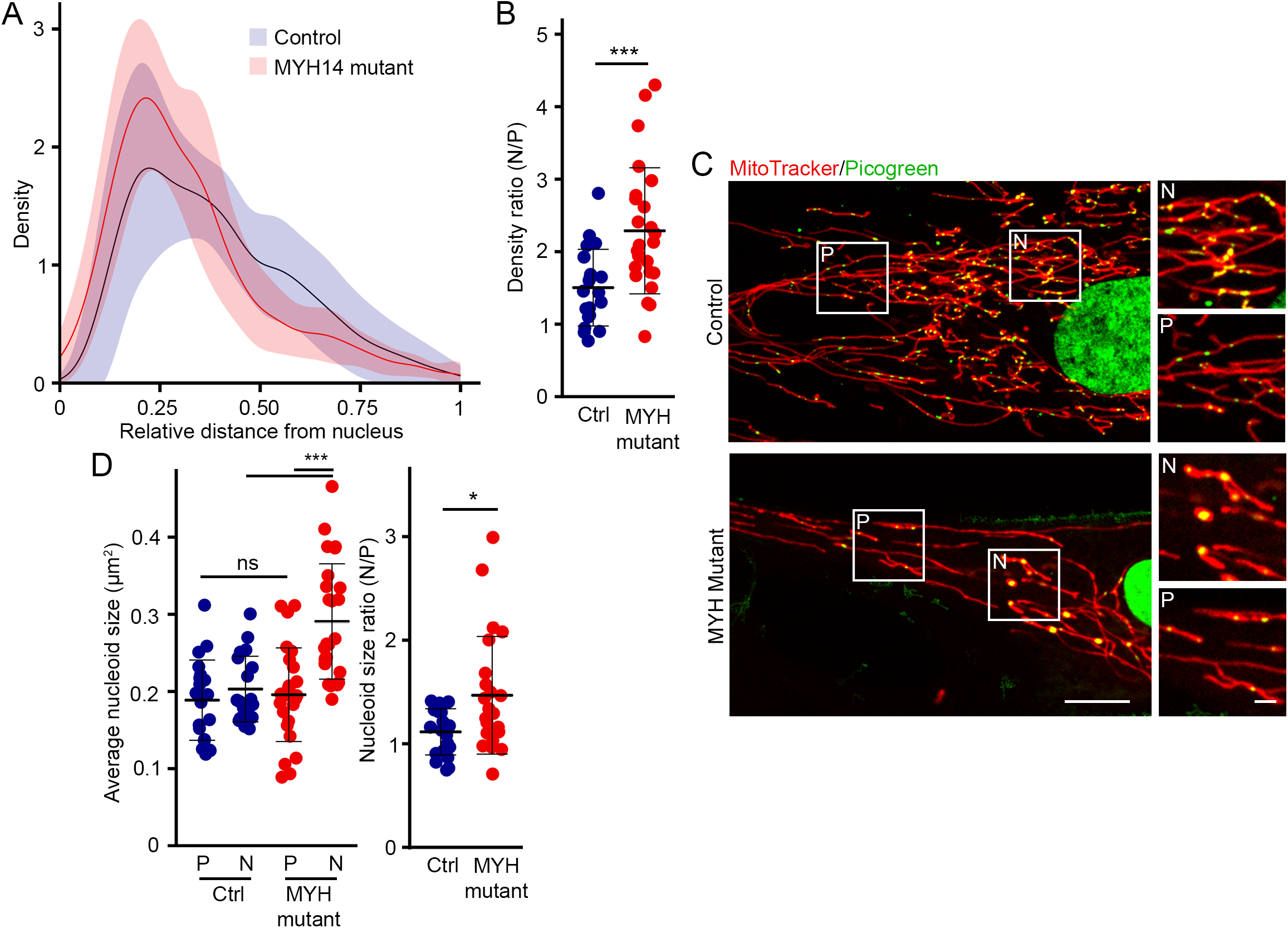
MYH14 mutation affects nucleoid spatial organization. (A) Density plot of nucleoids from human fibroblasts relative to the nucleus, where 0 corresponds to the nucleus and 1, the cell membrane. Solid lines represent the average distribution for MYH14 mutant primary fibroblasts (Red) and control fibroblasts (Blue, and the shaded areas, the SD (n=11 cells, MYH14 mutants; n=12 cells, Control). (B) Nucleoid density ratio (Perinuclear/Periphery) calculated from the number of nucleoids/μm mitochondria in each region. (C) Representative live cell confocal images of control and MYH14 mutant primary fibroblasts stained with mitotracker red (mitochondria) and picogreen (DNA) Scale bar, 10 μm; 2 μm for zoomed images. P, periphery; N, perinuclear. (D) Average nucleoid size in peripheral (P) and perinuclear (N) regions (Left) and nucleoid size ratio (Perinuclear/periphery) of control and MYH14 mutant primary fibroblasts. Each data point represents one cell. Bars represent the average of 20 cells ± SD. *, p<0.05, ***p<0.001, ns not significant. One-way ANOVA (Left), Two-tailed t-test (Right).

We previously reported that MYH14 mutant cells had fewer, but larger nucleoids than control cells (10). These were evident in mutant cells stained for nucleoids (picogreen) and mitochondria (mitotracker) (Fig. 6C). Importantly, these enlarged nucleoids (possibly cluster of mtDNAs) were restricted to the perinuclear region, which was confirmed by the quantification of nucleoid size close to the nucleus and in the periphery (average size and size ratio; Fig.6D). In contrast, control nucleoids were of similar size irrespective of their localisation (Fig. 6C-D). Thus, our results indicate that MYH14 significantly influences nucleoid maintenance and distribution, supporting the idea that mitochondrial fission is essential for the distribution of nucleoids.

### Dominant-negative mutation in DRP1 causes perinuclear nucleoid accumulation and altered nucleoid distribution

While these results support an important role for mitochondrial fission in the regulation of nucleoid distribution, it remained possible that the nucleoid phenotype we observed in MYH14 mutant cells was restricted to myosin defects. Thus, to confirm the role of fission in nucleoid distribution, we used primary fibroblasts from patients with a mutation in DRP1 (G362D), an essential component of the fission machinery (26). Mutations in DRP1 or its genetic deletion results in elongated and hyperconnected mitochondria and causes the formation of enlarged nucleoids termed mitobulbs (9, 26). While these previous studies did not determine the subcellular localisation of these mitobulbs, our results predict that they accumulate preferentially in the perinuclear region of mutant cells. To verify this, primary human fibroblast cells were stained for mitochondria (TMRM) and nucleoids (picogreen) and imaged by confocal microscopy (Fig. 7A). Similar to MYH14 mutant, the mitobulbs present in DRP1 mutants were mainly restricted to the perinuclear region of the cells (Fig. 7A). In fact, nucleoid size was significantly larger in the perinuclear region of the DRP1 mutants (Fig. 7B), and a larger fraction of total nucleoids was present in this region in mutant cells (perinuclear/periphery density ratio; Fig. 7C). Also, as with MYH14 mutants, the changes in nucleoid size and distribution were accompanied by a reduction in overall nucleoid density (Fig. 7D).

**Figure 7:**
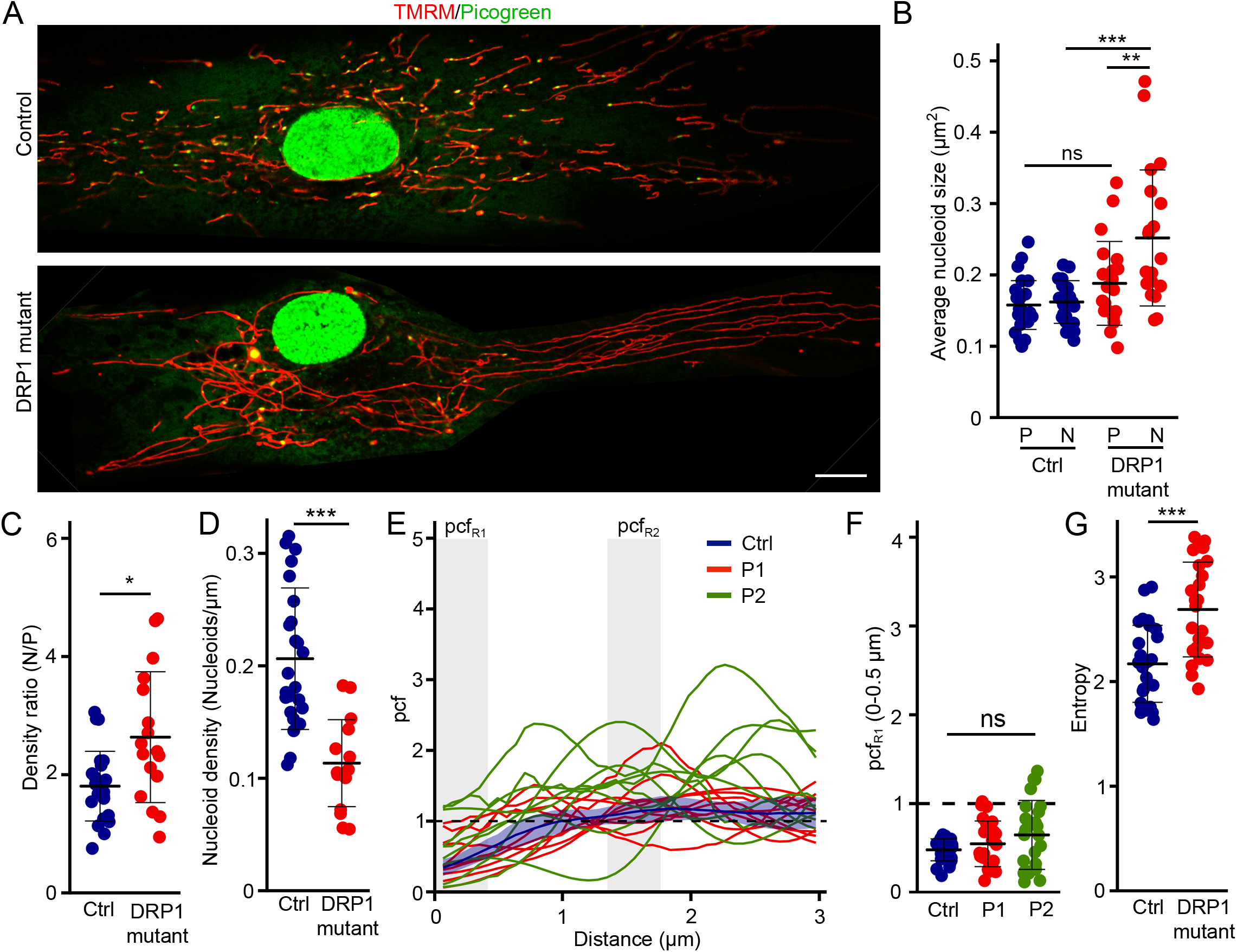
DRP1 is required for proper nucleoid distribution. (A) Representative live cell confocal images of control and DRP1 mutant primary fibroblasts stained with TMRM (mitochondria) and picogreen (DNA) Scale bar, 10 μm. (B) Average nucleoid size in peripheral (P) and perinuclear (N) regions of control and DRP1 mutant primary fibroblasts. Each data point represents one cell. Bars represent the average of 20 cells ± SD. **, p<0.01, ***p<0.001, ns not significant. One-way ANOVA. (C) Nucleoid density ratio (Perinuclear (N)/Periphery (P)) calculated from the number of nucleoids/μm mitochondria in each region. (D) Nucleoid density in control and DRP1 mutant primary fibroblasts. Each data point represents one cell. Bars represent the average of 20 cells ± SD. ***p<0.001. Two-tailed t-test. (E-G) Nucleoid distribution in control and DRP1 primary human fibroblasts. (E) pcf curves. The solid line for the control (Blue) represents the average distribution and the shaded area, the SD (n=9 cells). For the DRP1 mutants, each solid line represents one individual cell (n=9 cells per patient line), highlighting the variability of the pcf. The dashed line represents the expected random distribution and the grey areas, the distance for which the average pcf was quantified (F; pcfR1: 0-0.5μm). (F) Average pcf values at short distances. Each data point represents one cell. Bars represent the average of 30 cells ± SD. ns not significant. One-way ANOVA. (G) Entropy values calculated from the pcf curves in (E). Each data point represents one cell. Bars represent the average of 30 cells ± SD. ***p<0.001. Two-tailed t-test.

We then measured nucleoid distribution in DRP1 mutants. Similar to MYH14 mutant, the pcf of DRP1 mutant cells was variable, with some mutant cells showing greater correlation at short distances while others avoided each other at short distance (pcfR1-0-0.5μm) (Fig 7E-F). As with the MYH mutants, the change in pcf caused by DRP1 mutation also resulted in an increase in entropy (Fig. 7G). Overall, our data indicates that defects in mitochondrial fission impairs proper nucleoids distribution, resulting in their perinuclear clustering and enlargement.

### Synergistic effect of mitochondrial features influences nucleoid distribution in fission mutants

To understand how impaired mitochondrial fission leads to such alterations in nucleoid distribution, we first determined whether network features (mitochondrial length, connectivity) and nucleoid parameters (total nucleoids, size ratio) correlated with the pcf changes observed in the two fission mutants. To do this, we calculated Pearson coefficient between each parameter independently for each cell type. Control lines for MYH14 and DRP1 mutants behaved similarly, with the same parameters showing a correlation (Pearson coefficient ≥ ± 0.5, Boxed in Fig. 8A). Among these was a predictable correlation between network size and total nucleoid numbers, but also a correlation between these two parameters and entropy. Importantly, the correlation pattern clearly varied between genotypes (Fig. 8A), suggesting that each genotype behaves differently.

**Figure 8:**
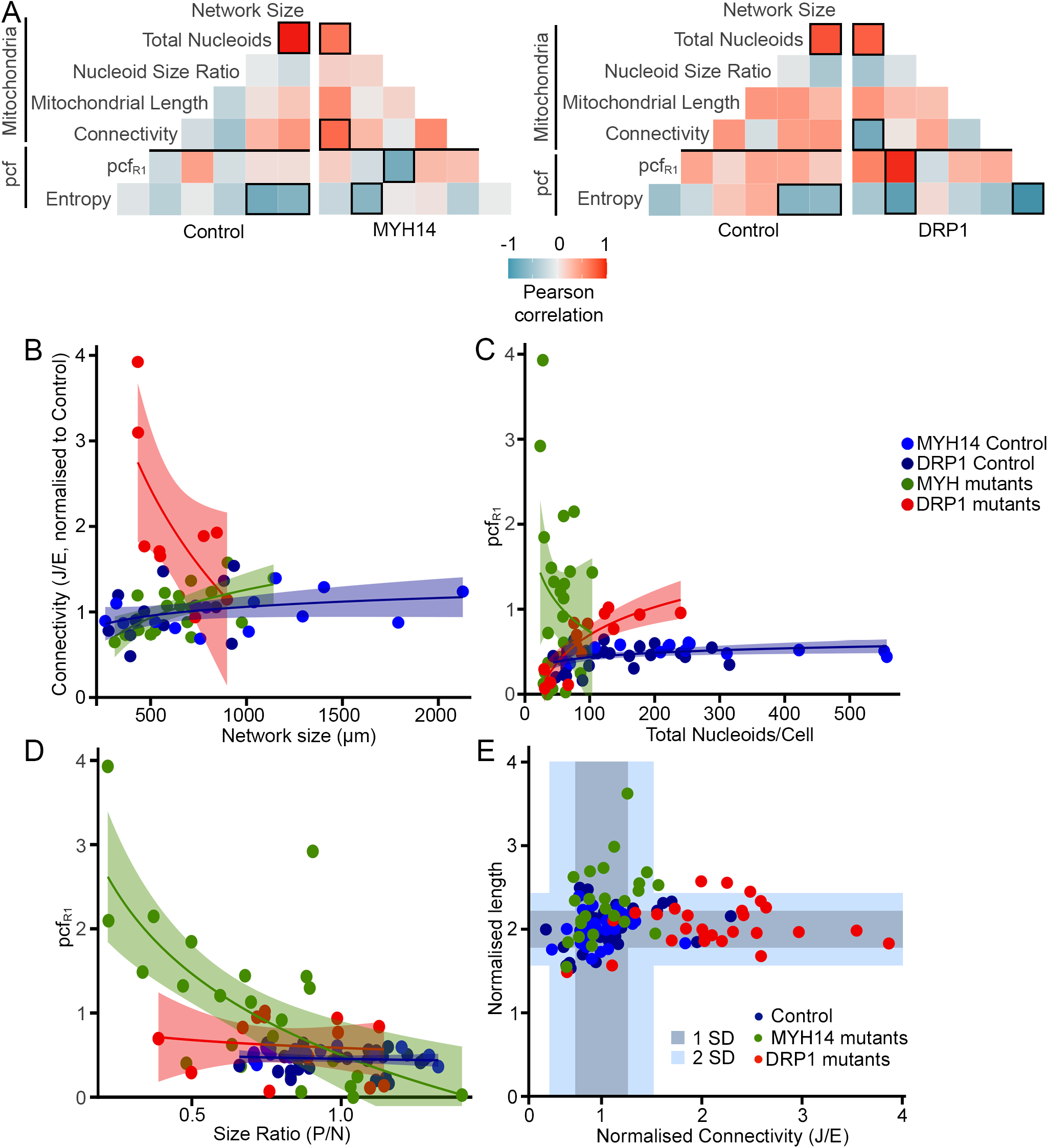
Fission mutants show a distinct relationship between mitochondrial parameters and the pcf. (A) Pearson correlation of pcf and mitochondrial parameters for the indicated genotypes. Number of cells used for the analysis: Control MYH14 (24), MYH14 mutants (25), Control DRP1 (17), DRP1 mutants (14) (B-D) Scatter plots showing the correlation between the indicated parameters for each genotype. The shaded areas represent the 95% confidence interval. Each data point represents one cell (n as in (A)). Linear regression formula y~log(x). (E) Distribution of mitochondrial connectivity and length across genotypes in individual cells. Connectivity was calculated as the number of junctions (J)/number of ends (E). For length, we used the number of mitochondria <2 μm but, to have an increasing value with increasing length, we used 1/this number. All values were normalized to the control for the same experiment. The shaded areas represent 1 SD and 2 SD from the control values. Each data point represents one cell.

The distinct behavior of each genotype was evident when comparing network features (network size vs connectivity shown in Fig. 8B; Pooled controls r^2^=0.07, p=0.10; MYH14 r^2^=0.32, p<0.01; DRP1 r^2^=0.38, p<0.05) but also when nucleoid parameters were correlated with the pcf. For example, the pcf at close distance (PcfR1) correlated with nucleoid number specifically in the DRP1 mutants (Fig. 8A, C; DRP1: r^2^=0.69, p<0.001; pooled controls: r^2^=0.16, p<0.01; MYH14: r^2^=0.00, p<=0.35), while pcfR1 correlated with nucleoid size ratio only in MYH14 mutants (Fig. 8A, D; pooled controls: r^2^=0.02, p=0.53; MYH14: r^2^=0.45, p<0.001; DRP1: r^2^=0.07, p=0.67). These results suggest that the differences observed across genotypes are the consequence of distinct changes in mitochondrial features. This is supported by the fact that mitochondrial networks were distinctly affected in MYH14 and DRP1 mutants: MYH14 mutation mainly caused mitochondrial elongation while DRP1 mutants showed a large increase in connectivity (Fig. 8E).

On the other hand, a principal component analysis (PCA) of the same genotypes indicated that both mutants segregated away from control cells (Fig. 9A), suggesting that the fission mutants nonetheless share common features relative to mitochondrial network features. In fact, examination of the correlation data indicated that the entropy was correlated with nucleoid content for all genotypes (Fig. 8A; Fig. 9B, overall: r^2^=0.49, p<0.001; pooled controls: r^2^=0.49, p<0.001; MYH14: r^2^=0.67, p<0.001; DRP1: r^2^=0.26, p<0.01), suggesting that pcf variability is a consequence of the smaller number of nucleoids present in the mutant cells (Fig 5A, 7D). In order to verify whether nucleoid density affects nucleoid distribution, we analyzed nucleoid distribution in control cells where nucleoid density was decreased to match that of MYH14 mutants. To do this, we modified Mitomate tracker to be able to randomly remove points from each input image and analyse it as if it was the actual image (which is distinct from Fig. 2E where several random distributions were averaged – see Methods). This resulted in alterations in pcf curves and entropy (Fig 9C-E) that were similar to those observed in MYH14 mutants (Fig. 5B-D), indicating that nucleoid density influences the nucleoid distribution pattern. Nevertheless, the relationship was different between control cells and MYH14 mutant cells (Fig. 9B, p<0.001), suggesting that factors other than nucleoid number affect the entropy. This is also supported by the observation that the relationship between the entropy and pcfR1 varied across genotypes (Fig. 9F; Pooled controls: r^2^=0.17, p<0.05; MYH14: r^2^=0.02, p=0.46; DRP1: r^2^=0.74, p<0.001).

**Figure 9:**
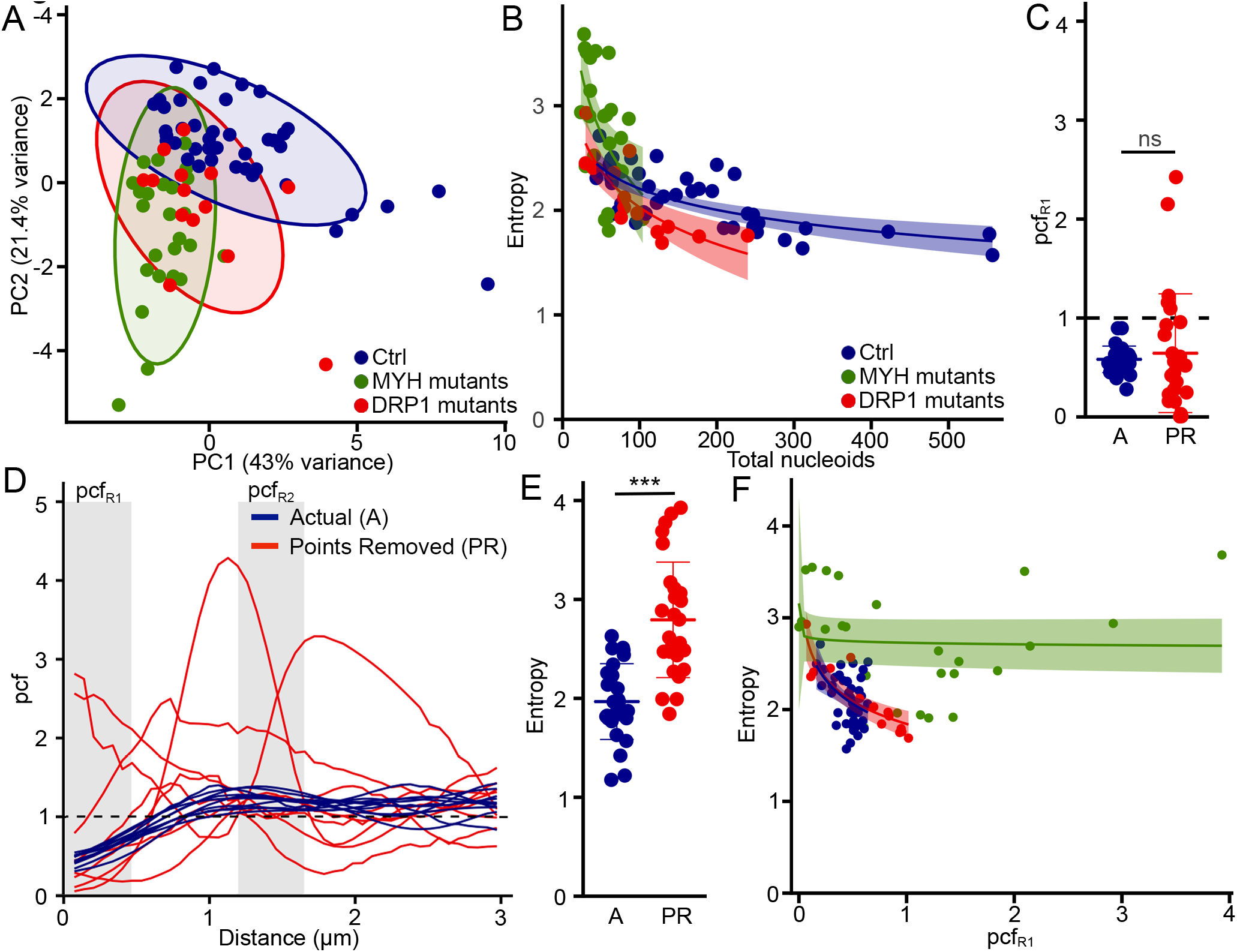
Nucleoid density influences the variability of the pcf curves (Entropy) across genotypes. (A) PCA analysis showing the segregation of mutant and control lines. Each data point represents one cell. Circles represent the 95% confidence interval (B) Dot plots showing the correlation between Entropy and total number of nucleoids for each genotype. The shaded areas represent the 95% confidence interval. Each data point represents one cell. (C-E) Decreasing nucleoid density in control cells recapitulates the pcf variability found in fission mutants. Individual cells are shown for pcfR1 values (C), pcf curves (D) and Entropy (E). (F) Relationship between pcfR1 and Entropy in different genotypes. The shaded areas represent the 95% confidence interval. Each data point represents one cell.

Altogether, these results suggest that while both fission mutants globally affect nucleoid features and their distribution in a similar way (Fig. 9A), the manner in which they modulate this process likely differs as a consequence of distinct changes in mitochondrial network features.

## Discussion

Mitochondrial function depends on the proper maintenance of mtDNA and its distribution across the mitochondrial networks. While mitochondrial fusion plays an important role in mtDNA maintenance (27), the role of mitochondrial fission in this process remains poorly understood. In fact, impaired fission is associated with the presence of enlarged nucleoids but not necessarily a loss of mtDNA content (9, 10, 12). In addition, while mitochondrial dynamics have been suggested to control nucleoid distribution across mitochondrial networks, the underlying mechanisms remain poorly understood. To address these questions, we developed an automated tool, Mitomate tracker to quantify nucleoid distribution within mitochondrial networks. Mitomate tracker allows to use minimally preprocessed images to provide detailed quantitative data on mitochondrial network and nucleoids features and distribution. We initially measured both nndist and pcf to quantify nucleoid distribution. However, we found that the pcf is a more robust metric than the nndist. This is likely because the pcf globally measures the distribution probability relative to an IRP instead of the single average distance provided by the nndist. In addition, unlike the nndist, the calculation of the pcf probability takes into account the fact that points are closer when density is higher, providing a normalization relative to density. While we used it here to demonstrate the importance of mitochondrial fission in the regulation of nucleoid distribution, it could be utilized to characterize the distribution of any mitochondrial proteins dispersed as individual foci along mitochondrial networks.

Our results indicate that in healthy cells, nucleoids are distributed in a semi-regular manner, with nucleoids strongly avoiding each other at closer distances (Fig 3A-B), consistent with previous manual nndist measurements in yeast and human cells (15, 16). In addition, we found that nucleoids have a distinctive spatial distribution pattern, with the perinuclear region having the highest nucleoid density relative to the size of the mitochondrial network. Nonetheless, the pcf (relative to an IRP) remained similar for both the perinuclear region and cell periphery (Fig 4D). This is consistent with a previous STED imaging study that measured absolute inter-nucleoid distances (not distances within the mitochondrial network as here) and found that while perinuclear nucleoids maintain a proper distance from their neighbours (nndist), the nndist increases with distance to the nucleus (14). Altogether, our results strongly support the notion that nucleoid distribution is actively regulated in the cells.

Recently, we have shown that mutation of the mitochondrial fission protein MYH14 reduced total nucleoid population without altering mtDNA content (10). Similarly, knock down or genetic ablation of DRP1 altered nucleoid content by causing the formation of mitobulbs (9, 12, 13). As these results suggest a potential role for fission in nucleoid maintenance, we have used both mutants to directly address the role of mitochondrial fission in the regulation of nucleoid distribution. Our results show that inhibiting mitochondrial fission disrupted nucleoid distribution as reflected by the high variability of the pcf curves and the perinuclear accumulation of enlarged nucleoids in both MYH14 and DRP1 mutants.

The fission mutants showed a notable variation in their nucleoid distribution, as seen by both pcf curves and their associated entropy. Specifically, some mutant cells showed strong short distance correlation while in others, nucleoids avoided each other. The reduced nucleoid density observed in MYH14 and DRP1 mutants significantly contributed to this variability (entropy). Nevertheless, the specific contribution of distinct nucleoid or network parameters to the pcf varied across genotypes, suggesting a complex interplay between nucleoid and mitochondria network topologies in regulating nucleoid distribution. Consistent with this, mitochondrial networks were distinctly affected in MYH14 and DRP1 mutants, likely reflecting the distinct role of these proteins in mitochondrial fission. While DRP1 is an essential fission protein required for the physical severing of mitochondrial tubules, MYH14 encodes one of three non-muscle myosin II proteins that are involved in the initial ER-mediated constriction of the mitochondrial tubule (10, 28–30). In addition, we previously found that the mitochondrial phenotype of MYH14 mutants was most evident in the peripheral area of the cells (10), which could further affect the nucleoid distribution pattern.

The fact that our findings differed between the MYH14 and DRP1 mutant cells is not surprising, given the disparate clinical phenotypes in the patients from whom they were isolated. The MYH14 patients developed axonal sensorimotor neuropathy and sensorineural hearing loss (10), whereas the DRP1 patient had severe central nervous system involvement (26). A question of interest for future study with this tool may be to determine whether it can resolve differences in nucleoid distribution correlated to phenotypic variation from mutations in the same gene, implicating differing mechanisms for alternate clinical presentations. For example, MFN2 is a major human disease gene associated with numerous mitochondrial functions (including mitochondrial fusion) and highly variable clinical phenotypes, which are not consistently correlated to cellular phenotypes (31). In the specific case of MYH14, other mutations result in isolated and severe hearing loss or later-onset sensorineural hearing loss (32–34). In the case of DRP1, phenotypes can vary, and recently severe cardiac involvement has been described from a novel mutation (35). The automated and quantitative approach described in this work presents an additional tool that may be valuable to genotypephenotype-mechanistic correlation.

It is nevertheless important to note that in our setup, mtDNA clusters less than about 300nm apart are not resolved and thus do not contribute to the pcf curves. While this probably does not affect wild-type cells, this could alter the analysis of the fission mutants as they contain enlarged nucleoids that likely represent a cluster of mtDNAs that failed to separate following mtDNA replication. It is thus possible that, under conditions where individual mtDNA molecules could be resolved, the pcf would detect a strong correlation at very short distances (< 300 nm) in these cells. Nevertheless, our data supports the idea that mitochondrial fission regulates nucleoid distribution and prevents nucleoid clustering to facilitate homogenous distribution of nucleoids within mitochondrial networks.

Another important finding of our study is that the accumulation of enlarged nucleoids in fission mutants is mainly restricted to the perinuclear region. As larger nucleoids likely contain a higher number of mtDNA copies, our results is consistent with an earlier report suggesting that mtDNA replication preferentially occurs in mitochondrial clusters near the nucleus (23). Mitochondrial fission could thus play an important role in nucleoid segregation and distribution by allowing newly synthesized nucleoid near the nucleus to spread to the peripheral mitochondrial network.

In conclusion, our results indicate that, while mitochondrial fission might not directly control mtDNA replication, it plays an essential role by regulating nucleoid distribution across mitochondrial networks. This process is likely required to facilitate homogenous distribution of mtDNA and OXPHOS protein subunits and its alteration in fission mutants likely contribute to the development of associated pathological conditions.

## Materials and Methods

### Reagents

Cell culture reagents were obtained from Wisent. Other chemicals were purchased from Sigma-Aldrich, except where indicated.

### Cell culture and live cell imaging

Primary human fibroblasts (controls, MYH14 mutants and DRP1 mutants) were generated from skin biopsies, collected as part of a research protocol, and written informed consent from participants was obtained (University of Calgary Research Ethics Board (MYH14 mutants), Research Ethics Board of the Children’s Hospital of Eastern Ontario (DRP1 mutants)). Biopsy samples were processed as described and cultured in Dulbecco’s modified Eagle’s medium (DMEM) containing 10% fetal bovine serum (FBS), supplemented with Penicillin/Streptomycin (100 IU/ml/100μL/mL)(26, 36). Immortalized Mouse Embryonic Fibroblasts (MEFs) were cultured in DMEM supplemented with 10% fetal bovine serum. For live cell imaging, cells were seeded onto glass bottom dishes and stained for 30 minutes with 250nM TMRM (Thermo fisher Scientific, T668) (MEFs and DRP1 mutant and control human fibroblasts) or 50nM Mitotracker Red (Thermo fisher scientific, M7512) (MYH14 mutant and control fibroblasts) and the DNA dye PicoGreen (Thermo Fisher Scientific, P11495) (3 μL/mL). After staining, cells were washed 3 times with pre-warmed 1X phosphate buffered saline (PBS), and normal growth media was added prior to imaging.

### Microscopy

Images for MEFs, DRP1 mutant fibroblasts and their wild-type control were acquired with a Leica TSC SP8 confocal microscope fitted with a 63x/1.40 oil objective using the optimal resolution for the wavelength (determined using the Leica software). Images from MYH14 cells and their control were taken with an Olympus spinning disc confocal system (Olympus SD-OSR) (UAPON 100XOTIRF/1·49 oil objective) operated by Metamorph software. The SD-OSR was equipped with a cellVivo incubation module to maintain cells at 37°C and 5% CO2 during live cell imaging.

### Image analysis using Mitomate tracker

Red and green channels were separated, nuclei were manually removed from the Picogreen channel and the images converted to 8-bit using ImageJ. The mitochondrial channel was then segmented using the ImageJ filter tubeness and global thresholding. To measure the correlation between nucleoids on the mitochondrial network, the images were analyzed using the R package spatstat (flowchart in Fig S2). This analysis requires two inputs: a linear network (mitochondria) and a point pattern (nucleoids). Mitochondrial components (tubules, ends (E), junctions (J)) were extracted from the segmented mitochondrial image using Momito (18) and used as the input for the linear network. Nucleoids were identified using the ImageJ plugin TrackMate (19) and the coordinates of the nucleoids associated with the mitochondrial network (within 6 pixels center to center) used to generate the point pattern. Overall 97±1% of the nucleoids were properly identified in control cells, although 38%of the larger nucleoids (>0.6 μm) present in mutant cells were identified as 2 or more nucleoids.

The pcf analysis was then carried using the spatstat *linearpcf* function for distances from 0 to 300 pixels with a bias correction at each end of the interval (*correction=“Ang”*) and a bandwidth of 5 pixels, while the nndist was calculated using the spatstat *nndist* function. As Spatstat computes the pcf for individual mitochondrial clusters, we had to sum the contribution of each cluster to generate the total pcf by taking into consideration the size of each mitochondrial cluster and the number of nucleoids that it contains. This was achieved as follows:

In Spatstat, the estimator for the pcf *ĝ_k_*(*r*) for a given subgraph *G_k_*(a mitochondrial cluster) is given by

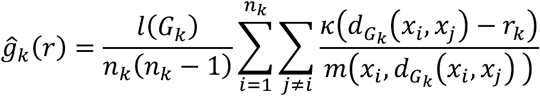

Where *k* is the gaussian kernel of 5 pixels used for smoothing and m is analogous to the perimeter for a network of radius *d_G_k__*(*x_i_, x_j_*) around the point *x_i_*. The length of the subgraph is *l*(*G_k_*) and it contain *n_k_* points. We have normalized the pcf *ĝ_k_*(*r*) for the whole network (graph, *G*) by

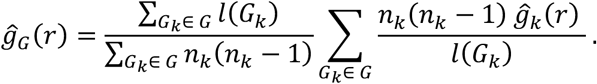

In addition, to avoid spurious effects caused by variation in nucleoid density and network features (length and connectivity), we normalised both nndist and pcf by dividing the observed value (actual nndist or pcf) by the value obtained using an IRP with the same point density distributed across the same network. For the pcf, this was done for each distance (r) measured. The randomised point pattern used to correct for network effects was generated using the spatstat *runiflpp* function with the same number of points as the actual point pattern. Each image was run six times and averaged. To simplify the process, the analysis was automated using a Java script run on Eclipse. Network features and the total number of nucleoids were directly extracted from the Mitomate tracker analysis, except for the connectivity that was defined as the total number of junctions (J)/ total number of mitochondrial ends (E) for each individual cell (18).

To compare the effect of nucleoid and mitochondrial features on the pcf with random distributions, we generated random point patterns with a similar density as that of the actual point patterns (distinct from the IRP used above to correct for network effects) using the same Java script. However, as each of these IRPs represent a specific distribution that can somewhat vary from the expected random distribution (especially when point density is low), 6 distributions were averaged for each experiment to avoid measuring effects due to specific random distributions. In the case of experiments where nucleoids were randomly removed, an individual distribution with points removed was considered as the actual data that was normalised over the average of 6 random distributions.

### Analysis of spatial distribution

8-bit nucleoid images and segmented mitochondria networks were manually separated into perinuclear and peripheral mitochondrial clusters using ImageJ. The perinuclear area was defined as the region within 23 μm from the nuclear membrane, although care was taken to keep individual mitochondrial clusters intact when separating the mitochondrial network (each independent cluster were labeled as either perinuclear or peripheral). These images were then used to measure nucleoid size (ImageJ - *Analyse Particle* function) and region-specific pcfs (Mitomate tracker). Density distributions were calculated by first determining the coordinates of each nucleoid relative to the nucleus using ImageJ (*Analyse Particle* function) and compiling the result using the R function *density*.

### Data analysis and statistics

All data analysis was done in R. To quantify the entropy, pcf values for each distance first needed to be converted into a character string. This was achieved by first converting the decimal numbers into a whole number between 1 and 26 and attributing a letter to each number. The entropy of the resulting character string was then calculated using the *entropy* function of the R package acss (37). The PCA was done using the R function *prcomp* while Pearson correlations were determined using the *ggcorr* function (GGally package).

Statistical analysis was done using Student’s t test (between 2 groups) or one-way ANOVA with a tukey post hoc test (multiple comparisons). Differences between nucleoid distributions were calculated using a KS test (*ks.test* function from the stats package). Linear regressions for correlation analysis were calculated using a lm method with y~log(x) as the general formula.

## Acknowledgements

We thank Andrea Bertolo for his help with the entropy calculations and A Micheil Innes who, with GP, provided the MYH14 mutants. This work was supported by grants from the Natural Sciences and Engineering Research Council of Canada and the Fondation de l’UQTR to MG, as well as the Alberta Children’s Hospital Research Institute to TS. HSI was supported by a Queen Elizabeth II Diamond Jubilee Scholarship and a FRQ-NT scholarship, while RS was supported by a QEII Graduate Scholarship. MO received a CIHR undergraduate scholarship for this work.

## Author contribution

HSI and RS performed the cell biology experiments. MO designed and tested the analysis tools, and did the experimental analysis using them. HSI, MO and MG designed the experiments. HSI, MO, JDG and MG analysed the data. RS and TES provided the MYH14 raw data. MAL and GP provided the clinical samples. HSI and MG wrote the paper. All authors reviewed and discussed the manuscript.

---

**Supplementary figure 1:**
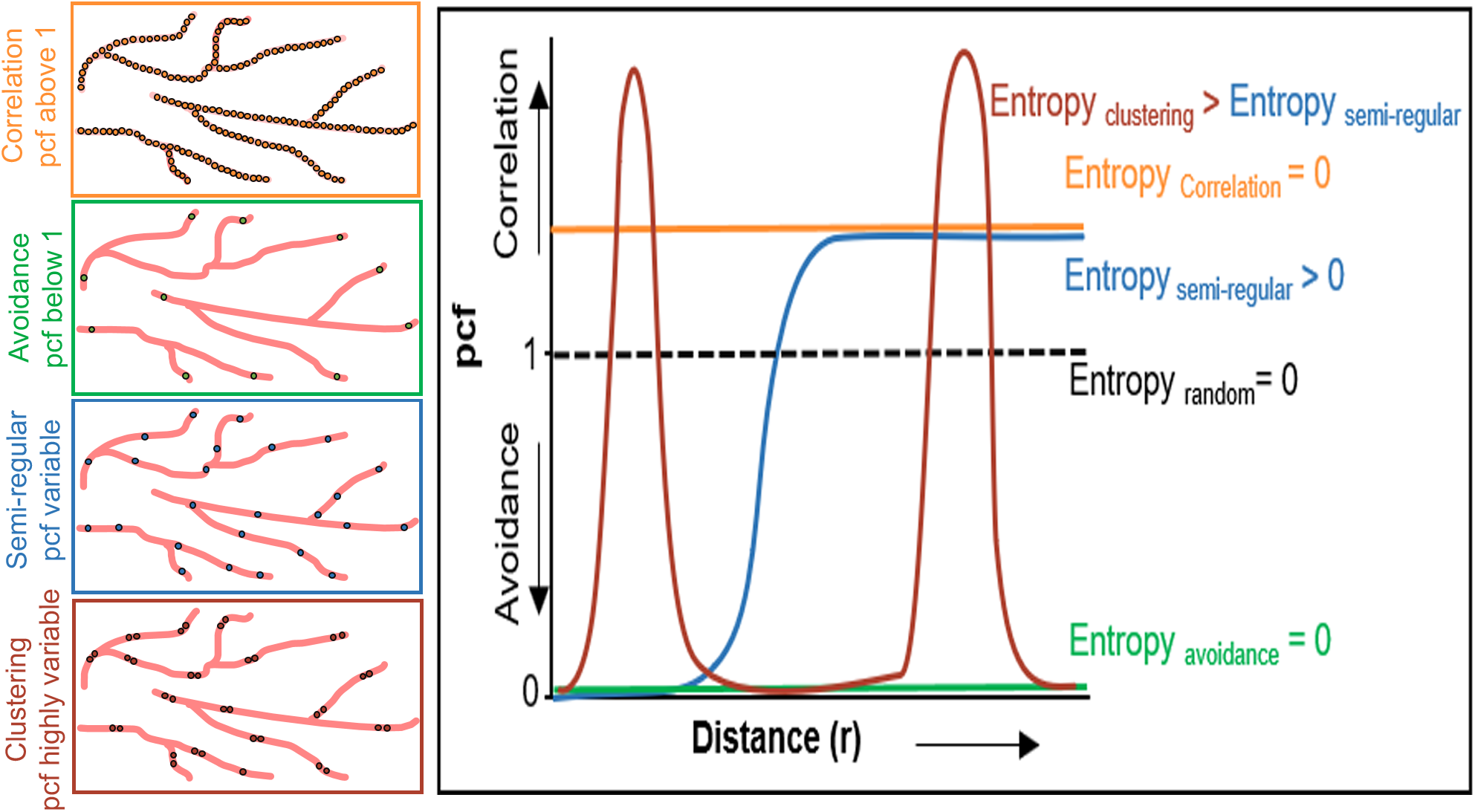
Entropy, a measure of variability in nucleoid distribution. The left panel represent different possible nucleoid distributions within mitochondrial networks. The right panel shows the expected pcf curves for the same distributions. Pcf value above 1 suggest correlation and anything below 1 suggest avoidance between nucleoids. The entropy, calculated from the pcf curves, increases with increasing variability of the pcf curve (an horizontal line has an entropy value of zero).

**Supplementary figure 2:**
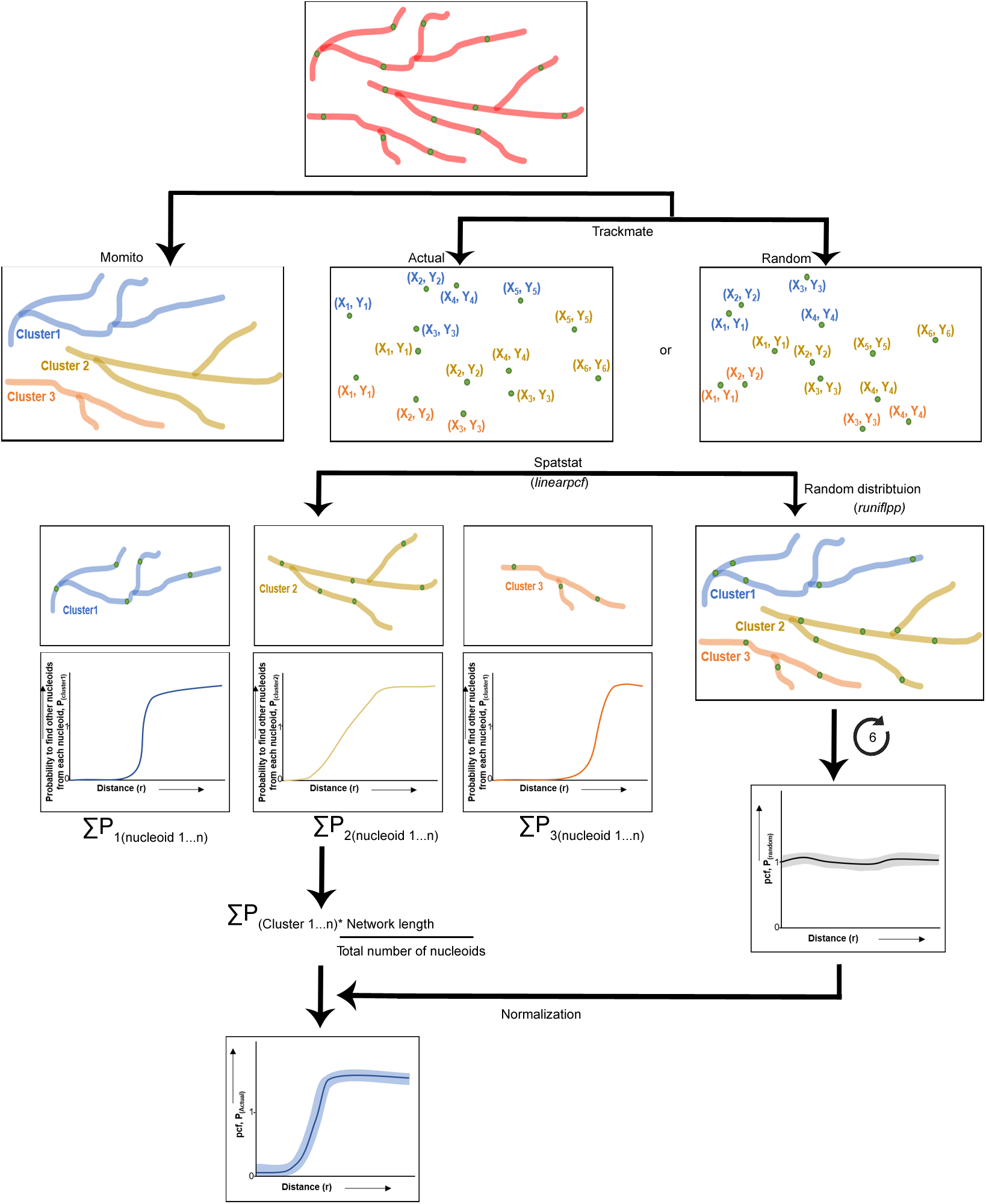
Workflow of Mitomate tracker. The preprocessed mitochondrial images are analyzed by Momito to extract mitochondrial network features. The coordinates of nucleoid in actual or random distributions are extracted by the Image J plugin Trackmate. Based on this information, Spatstat measures the probability of nucleoid distribution in each individual mitochondrial cluster (independent mitochondrial network). Overall nucleoid distribution is calculated by summing up the probability in all mitochondrial clusters. To take into account the effect of network features and nucleoid density, 6 random distributions (using runiflpp function) are averaged and used to normalize the actual distribution.

## References

1. T. Sasaki, Y. Sato, T. Higashiyama, N. Sasaki, Live imaging reveals the dynamics and regulation of mitochondrial nucleoids during the cell cycle in Fucci2-HeLa cells. Sci Rep 7, 11257 (2017).

2. M. Giacomello, A. Pyakurel, C. Glytsou, L. Scorrano, The cell biology of mitochondrial membrane dynamics. Nature reviews. Molecular cell biology 21, 204–224 (2020).

3. G. S. Barsh et al., Mitochondrial fusion is required for regulation of mitochondrial DNA replication. PLOS Genetics 15 (2019).

4. A. W. El-Hattab, W. J. Craigen, L. J. C. Wong, F. Scaglia, “Mitochondrial DNA Maintenance Defects Overview” in GeneReviews((R)), M. P. Adam et al., Eds. (Seattle (WA), 1993).

5. C. Viscomi, M. Zeviani, MtDNA-maintenance defects: syndromes and genes. Journal of inherited metabolic disease 40, 587–599 (2017).

6. J. R. Friedman et al., ER tubules mark sites of mitochondrial division. Science 334, 358–362 (2011).

7. A. Pagliuso, P. Cossart, F. Stavru, The ever-growing complexity of the mitochondrial fission machinery. Cellular and molecular life sciences: CMLS 75, 355–374 (2018).

8. S. C. Lewis, L. F. Uchiyama, J. Nunnari, ER-mitochondria contacts couple mtDNA synthesis with mitochondrial division in human cells. Science 353, aaf5549 (2016).

9. R. Ban-Ishihara, T. Ishihara, N. Sasaki, K. Mihara, N. Ishihara, Dynamics of nucleoid structure regulated by mitochondrial fission contributes to cristae reformation and release of cytochrome c. Proc Natl Acad Sci U S A 110, 11863–11868 (2013).

10. W. Almutawa et al., The R941L mutation in MYH14 disrupts mitochondrial fission and associates with peripheral neuropathy. EBioMedicine 45, 379–392 (2019).

11. P. A. Parone et al., Preventing Mitochondrial Fission Impairs Mitochondrial Function and Leads to Loss of Mitochondrial DNA. PLoS ONE 3 (2008).

12. A. Ota, T. Ishihara, N. Ishihara, Mitochondrial nucleoid morphology and respiratory function are altered in Drp1-deficient HeLa cells. Journal of biochemistry 167, 287–294 (2020).

13. T. Ishihara et al., Dynamics of mitochondrial DNA nucleoids regulated by mitochondrial fission is essential for maintenance of homogeneously active mitochondria during neonatal heart development. Mol Cell Biol 35, 211–223 (2015).

14. C. Kukat et al., Super-resolution microscopy reveals that mammalian mitochondrial nucleoids have a uniform size and frequently contain a single copy of mtDNA. Proc Natl Acad Sci U S A 108, 13534–13539 (2011).

15. J. Tauber et al., Distribution of mitochondrial nucleoids upon mitochondrial network fragmentation and network reintegration in HEPG2 cells. Int J Biochem Cell Biol 45, 593–603 (2013).

16. C. Osman, T. R. Noriega, V. Okreglak, J. C. Fung, P. Walter, Integrity of the yeast mitochondrial genome, but not its distribution and inheritance, relies on mitochondrial fission and fusion. Proc Natl Acad Sci U S A 112, E947–956 (2015).

17. R. Jajoo et al., Accurate concentration control of mitochondria and nucleoids. Science 351, 169–172 (2016).

18. M. Ouellet, G. Guillebaud, V. Gervais, D. Lupien St-Pierre, M. Germain, A novel algorithm identifies stress-induced alterations in mitochondrial connectivity and inner membrane structure from confocal images. PLOS Computational Biology 13 (2017).

19. J. Y. Tinevez et al., TrackMate: An open and extensible platform for single-particle tracking. Methods 115, 80–90 (2017).

20. A. Baddeley, E. Rubek, R. Turner, Spatial Point Patterns: Methodology and Applications with R (Chapman and Hall/CRC Press, London, 2015).

21. C. E. Shannon, A Mathematical Theory of Communication. Bell System Technical Journal 27, 623–656 (1948).

22. C. E. Shannon, A Mathematical Theory of Communication. Bell System Technical Journal 27, 379–423 (1948).

23. A. F. Davis, D. A. Clayton, In situ localization of mitochondrial DNA replication in intact mammalian cells. J Cell Biol 135, 883–893 (1996).

24. T. J. Collins, Mitochondria are morphologically and functionally heterogeneous within cells. The EMBO Journal 21, 1616–1627 (2002).

25. C. Wang et al., Dynamic tubulation of mitochondria drives mitochondrial network formation. Cell Research 25, 1108–1120 (2015).

26. J. R. Vanstone et al., DNM1L-related mitochondrial fission defect presenting as refractory epilepsy. European journal of human genetics: EJHG 24, 1084–1088 (2016).

27. E. Silva Ramos et al., Mitochondrial fusion is required for regulation of mitochondrial DNA replication. PLoS Genet 15, e1008085 (2019).

28. S. C. Kamerkar, F. Kraus, A. J. Sharpe, T. J. Pucadyil, M. T. Ryan, Dynamin-related protein 1 has membrane constricting and severing abilities sufficient for mitochondrial and peroxisomal fission. Nature Communications 9 (2018).

29. E. Smirnova, D.-L. Shurland, S. N. Ryazantsev, A. M. van der Bliek, A Human Dynamin-related Protein Controls the Distribution of Mitochondria. Journal of Cell Biology 143, 351–358 (1998).

30. F. Korobova, Timothy J. Gauvin, Henry N. Higgs, A Role for Myosin II in Mammalian Mitochondrial Fission. Current Biology 24, 409–414 (2014).

31. G. Sharma et al., Characterization of a novel variant in the HR1 domain of MFN2 in a patient with ataxia, optic atrophy and sensorineural hearing loss. 10.1101/2021.01.11.426268 (2021).

32. B. J. Kim et al., Discovery of MYH14 as an important and unique deafness gene causing prelingually severe autosomal dominant nonsyndromic hearing loss. J Gene Med 19 (2017).

33. M. Wang et al., A novel MYH14 mutation in a Chinese family with autosomal dominant nonsyndromic hearing loss. BMC medical genetics 21, 154 (2020).

34. F. Donaudy et al., Nonmuscle myosin heavy-chain gene MYH14 is expressed in cochlea and mutated in patients affected by autosomal dominant hearing impairment (DFNA4). American journal of human genetics 74, 770–776 (2004).

35. D. Vandeleur et al., Novel and lethal case of cardiac involvement in DNM1L mitochondrial encephalopathy. American journal of medical genetics. Part A 179, 2486–2489 (2019).

36. K. Martens et al., Case Report: Calpainopathy Presenting After Bone Marrow Transplantation, With Studies of Donor Genetic Content in Various Tissue Types. Frontiers in neurology 11, 604547 (2020).

37. H. S. N. Gauvrit, F. Soler-Toscano, H. Zenil, Algorithmic complexity for psychology. A user-friendly implementation of the coding theorem method. arXiv 1409.4080 (2014).

